# Kawasaki Disease patient stratification and pathway analysis based on host transcriptomic and proteomic profiles

**DOI:** 10.1101/2021.03.18.435948

**Authors:** Heather Jackson, Stephanie Menikou, Shea Hamilton, Andrew McArdle, Chisato Shimizu, Rachel Galassini, Honglei Huang, Jihoon Kim, Adriana Tremoulet, Marien de Jonge, Taco Kuijpers, Victoria Wright, Jane Burns, Climent Casals-Pascual, Jethro Herberg, Mike Levin, Myrsini Kaforou on behalf of the PERFORM consortium

**Affiliations:** Faculty of Medicine, Imperial College London, London, SW7 2AZ; Health Sciences, University of California San Diego, La Jolla, CA 92093; Oxford Biomedica (UK) Ltd, Oxford, OX4 6LT; Radboud University Medical Center, Nijmegen, 6525; Amsterdam University Medical Center (AMC), Amsterdam, 1105; Wellcome Trust Centre for Human Genetics, University of Oxford, Oxford, OX3 7BN

**Keywords:** infectious diseases, paediatrics, transcriptomics, proteomics, Kawasaki Disease, host ‘omics, systems biology, pathway analysis, clustering, classification

## Abstract

The aetiology of Kawasaki Disease (KD), an acute inflammatory disorder of childhood, remains unknown despite various triggers of KD having been proposed. Host ‘omic profiles offer insights into the host response to infection and inflammation, with the interrogation of multiple ‘omic levels in parallel providing a more comprehensive picture. We used differential abundance analysis, pathway analysis, clustering and classification techniques to explore whether the host response in KD is more similar to the response to bacterial or viral infection at the transcriptomic and proteomic levels through comparison of ‘omic profiles from children with KD to those with bacterial and viral infections. Pathways activated in patients with KD included those involved in anti-viral and anti-bacterial responses. Unsupervised clustering showed that the majority of KD patients clustered with bacterial patients on both ‘omic levels, whilst application of diagnostic signatures specific for bacterial and viral infections revealed that many transcriptomic KD samples had low probabilities of having bacterial or viral infections, suggesting that KD may be triggered by a different process not typical of either common bacterial or viral infections. Clustering based on the transcriptomic and proteomic responses during KD revealed three clusters of KD patients on both ‘omic levels, suggesting heterogeneity within the inflammatory response during KD. The observed heterogeneity may reflect differences in the host response to a common trigger, or variation dependent on different triggers of the condition.

## 1. Introduction

Kawasaki disease (KD) is an acute inflammatory disorder first described in Japan over 50 years ago [1]. KD occurs most frequently in children under five years of age [2]. Untreated KD leads to the formation of coronary artery aneurysms (CAAs) in 10-30% of children [3–5], causing it to be the most common cause of acquired heart disease in children in Europe, Japan and North America [6].

The aetiology of KD remains unknown, however the seasonality and epidemicity seen in areas of high incidence, including Japan, suggest that it is caused by an infectious trigger [7]. The current consensus is that, in some genetically predisposed children, an infectious trigger initiates an abnormal immune response [8,9]. Multiple viral and bacterial pathogens have been suggested as candidates for the trigger, in addition to airborne environmental and fungal triggers [8,10]. Despite the many theories postulated, none have been independently confirmed, and some studies have concluded that KD is likely to be caused by multiple environmental triggers [11].

As the coronavirus disease 2019 (COVID-19) pandemic evolved in early to mid-2020, an increase in cases of children with unusual febrile illnesses, some with features resembling KD, was observed [12]. This new condition, which was later termed “Paediatric Inflammatory Multisystem Syndrome Temporally associated with SARS-CoV-2”, or “Multisystem Inflammatory Syndrome in Children” (PIMS-TS or MIS-C) [12–15], tends to arise several weeks after SARS-CoV-2 infection [14]. The finding of increased KD-like cases after the emergence of a novel viral pathogen raises questions about whether more than one type of trigger might cause KD, and whether KD might represent a constellation of overlapping inflammatory syndromes.

Study of host transcriptomic and proteomic profiles can reveal perturbations caused by infection or inflammation. Comparison of the transcriptional response in different diseases has revealed different host responses to individual pathogens such as TB, bacterial and viral infections [16,17]. Previous studies of host ‘omics in the context of KD have characterised the pathways involved in the disease and have established biomarker signatures with diagnostic potential [18,19]. Interrogating multiple ‘omic datasets in parallel provides more accurate insights into the molecular dynamics of infection as information captured in one ‘omic layer might not necessarily be captured in another ‘omic layer.

We explored the host transcriptomic and proteomic profiles of children with KD and those with viral and bacterial infections, aiming to elucidate whether the inflammatory response in KD is more similar to that of a bacterial or viral infection, or indeed neither. We also used the approach to explore the heterogeneity within the transcriptional and translational response of patients with KD.

## 2. Results

### 2.1. Description of datasets

Whole blood transcriptomic profiles generated from 414 children were included in the analysis, obtained from children with Kawasaki Disease (KD; n = 178), confirmed (definite) bacterial infection (DB; n = 54), confirmed (definite) viral infection (DV; n = 120), and healthy controls (HC; n = 62). Two transcriptomic datasets were used. The ‘discovery’ transcriptomic dataset, which was generated by HumanHT-12 version 4.0 BeadChips, was used for all steps of the analysis. The ‘validation’ transcriptomic dataset, which was created by merging two datasets generated by HumanHT-12 version 3.0 and 4.0 BeadChips, was used to test the classifiers trained on the discovery dataset.

In addition, proteomic profiles from the plasma or serum of 329 children in the same groups were studied: from children with KD (n = 52), DB (n = 121) and DV (n = 106) infections, and HC (n = 50). Liquid chromatography with tandem mass spectrometry (LC-MS/MS) and the SomaScan [20] platform were used to generate the proteomic ‘discovery’ and ‘validation’ datasets, respectively. The ‘discovery’ proteomic dataset, generated from plasma samples using LC-MS/MS, was used for all steps of the analysis. The ‘validation’ proteomic dataset, generated from serum samples using the SomaScan platform [20], was used to test the classifiers trained on the discovery dataset.

On both ‘omic levels, the datasets that were used as ‘discovery’ datasets were selected due to their higher number of bacterial and viral samples. There was no overlap between the patients included in the proteomic datasets and those included in the transcriptomic datasets.

KD patients were defined according to AHA guidelines [21]. DB patients had a bacterial pathogen identified in a sample from a sterile site. DV patients had a virus identified that was consistent with the presenting syndrome; had no bacteria identified in blood or relevant culture sites; and had a CRP of <60mg/L. Further details on the clinical definitions used to define the DV and DB groups can be found in the Supplementary Text.

The median ages (months) of KD patients in the transcriptomic discovery and validation datasets were 26 (IQR: 29) and 37 (IQR: 34), respectively. The proportions of male KD patients were 55% and 60% for the transcriptomic discovery and validation datasets, respectively. For the proteomic KD group, the median ages (months) were 30 (IQR: 36) and 16 (IQR: 39) for the discovery and validation datasets, respectively. The proportion of males was 69% for both the discovery and validation datasets (Table S1). Table S2 contains clinical information for the KD patients included in the four datasets analysed. The causative pathogens for the patients with bacterial and viral infections from all datasets are shown in Table S3. The median duration of fever when the blood sample was taken for transcriptomic analysis from KD patients was 5 (range of 2-7 days) and 6 days (range of 2-10 days) for the discovery and validation datasets, respectively. For the proteomic KD samples, the median duration of fever when the sample was taken was 7 (range of 3-20 days) and 6.5 days (range of 4-22 days) for the discovery and validation dataset, respectively.

**Table 1.**
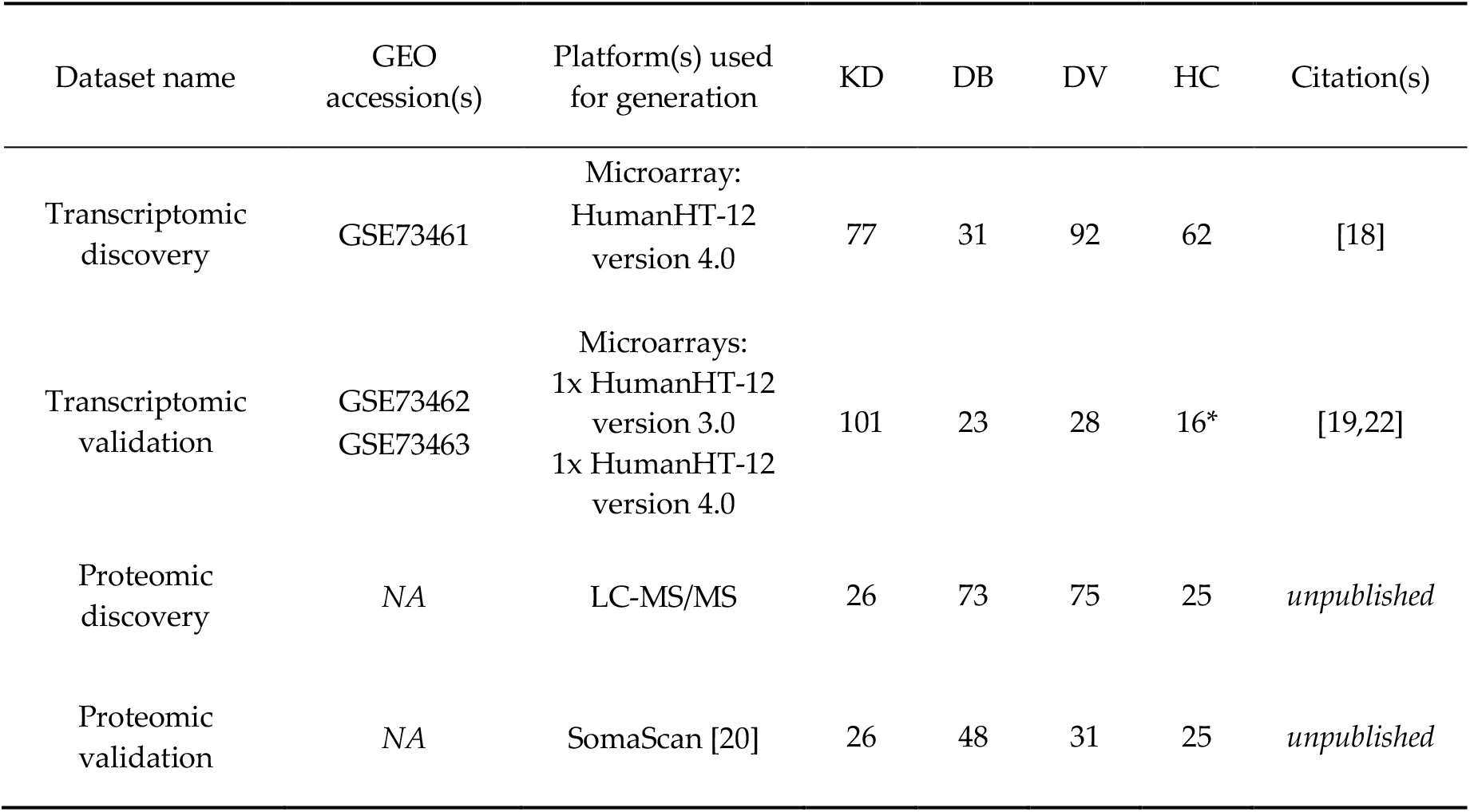
The datasets used in the analysis. KD, DB, DV and HC are abbreviations for Kawasaki Disease, definite bacterial, definite viral, and healthy control, respectively. LC-MS/MS is an abbreviation for liquid chromatography with tandem mass spectrometry. * = not used in analysis.

| Dataset name | GEO accession(s) | Platform(s) used for generation | KD | DB | DV | HC | Citation(s) |
| --- | --- | --- | --- | --- | --- | --- | --- |
| Transcriptomic discovery | GSE73461 | Microarray:<br>HumanHT-12<br>version 4.0 | 77 | 31 | 92 | 62 | [18] |
| Transcriptomic validation | GSE73462<br>GSE73463 | Microarrays:<br>1x HumanHT-12<br>version 3.0<br>1x HumanHT-12<br>version 4.0 | 101 | 23 | 28 | 16* | [19,22] |
| Proteomic discovery | NA | LC-MS/MS | 26 | 73 | 75 | 25 | *unpublished* |
| Proteomic validation | NA | SomaScan [20] | 26 | 48 | 31 | 25 | *unpublished* |

### 2.2. Comparison of Kawasaki Disease to Bacterial and Viral infection

We explored whether the host response during KD is more similar to the host response during bacterial or viral infections using transcriptomic (gene-level) and proteomic data. We first assessed the variance in the discovery datasets using Principal Component Analysis (PCA; Fig. S1-S2). In the transcriptomic dataset, PC1 (29.24%) appears to be capturing lymphocyte number and disease group, with the KD patients located between the bacterial and viral groups. In the proteomic dataset, PC1 (29.18%) appears to be capturing variation caused by age differences, while PC2 (13.39%) and PC3 (10.56%) strongly capture the disease group effects, with the KD patients grouped together between the clearly separated bacterial and viral groups.

#### 2.2.1. Differential abundance analysis

Limma [23] was used to identify genes and proteins differentially abundant between each disease group (KD, DB, DV) and healthy controls (HC), whilst accounting for age, sex and, for the transcriptomic dataset, immune cell proportions. Features were considered significantly differentially abundant (SDA) at a FDR of 5%. Differential abundance analysis was applied to 13035 genes and 344 proteins. For the transcriptomics, 3,218, 3,124, and 4,663 genes were SDA between KD vs HC, DB vs HC, and DV vs HC, respectively. For the proteomics, 113, 125, and 78 proteins were SDA between KD vs HC, DB vs HC, and DV vs HC, respectively. Genes and proteins SDA between KD vs HC are listed in the Supplementary File 1.

#### 2.2.2. Pathway analysis

The lists of SDA features identified in *2.2.1* were subjected to pathway analysis using g:Profiler2 [24] to determine which pathways were upregulated and downregulated in the three disease groups in the discovery datasets for the transcriptomic (Fig.1a) and proteomic (Fig. 1b) datasets. The full lists of pathways are provided in Supplementary File 2.

**Figure 1a.**
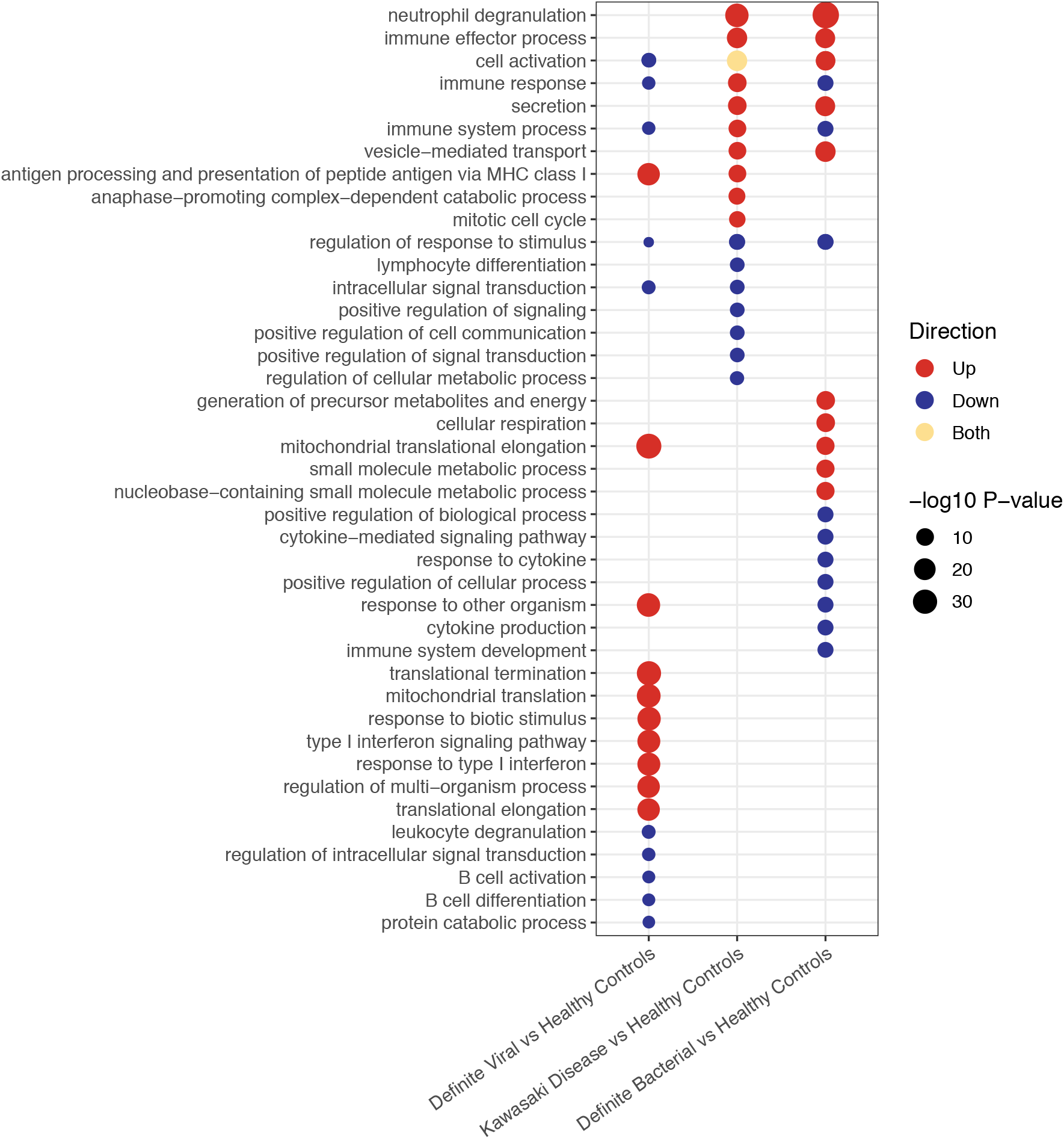
Pathways upregulated and downregulated in bacterial, KD and viral patients compared to healthy controls in the transcriptomic dataset.

**Figure 1b.**
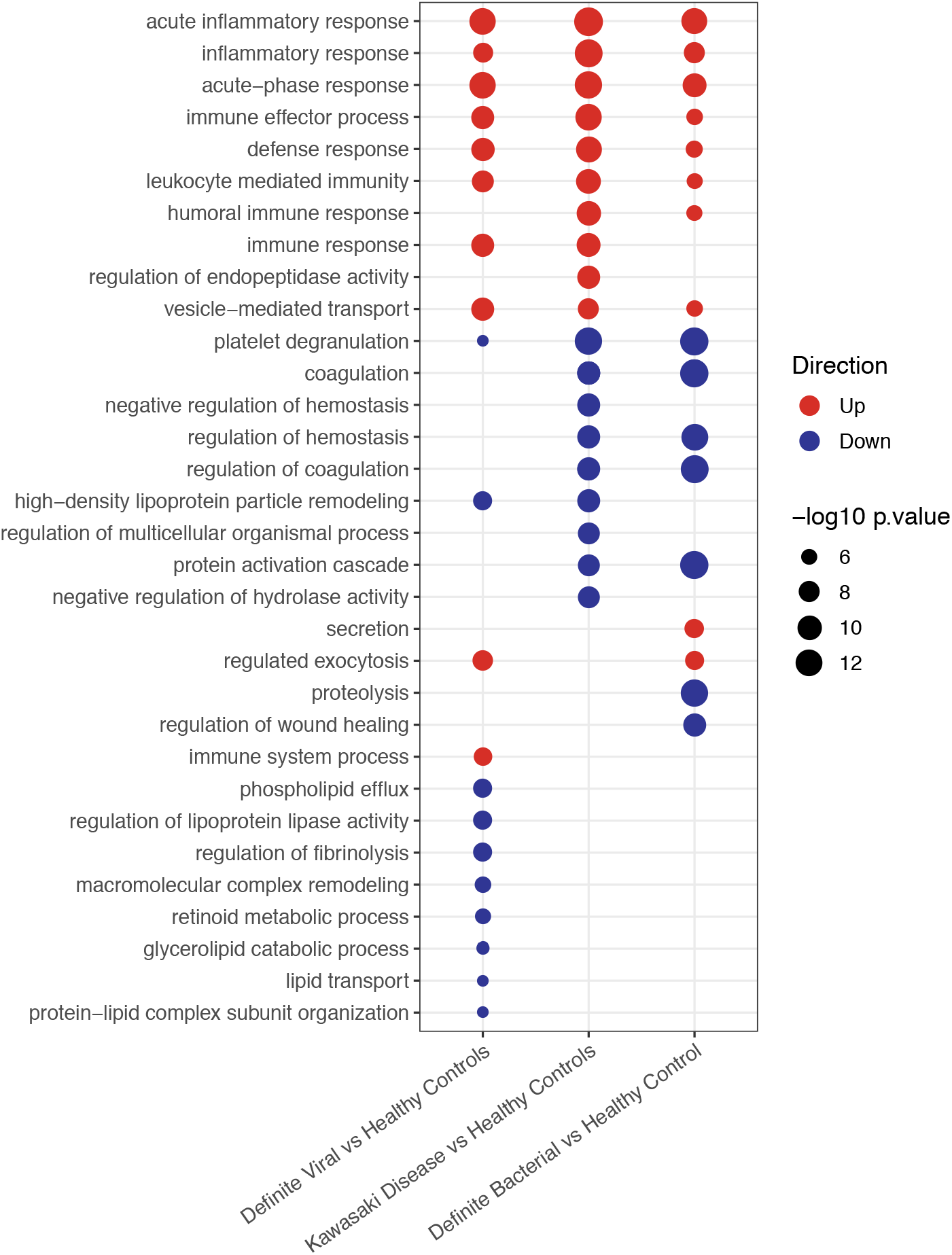
Pathways upregulated and downregulated in bacterial, KD and viral patients compared to healthy controls in the proteomic dataset.

In the transcriptomic pathway analysis, some pathways were found to be enriched across 2 or 3 of the disease conditions, whereas others were found in a single condition (Fig. 1a). For example, neutrophil degranulation, which was the top pathway in both KD and bacterial infections, and vesicle-mediated transport were both upregulated in KD and bacterial infections, whereas antigen presentation via MHC class I was upregulated in KD and viral infections. Of the top 17 pathways enriched in KD, 6 pathways were also present with concordant directions in the top bacterial pathways and 3 were present with concordant directions in the top viral pathways. Seven were unique to KD.

In the proteomics data, all pathways overlapping between KD and either bacterial or viral infections had concordant directions of regulation (Fig. 1b). Of the top 19 pathways enriched in KD, 13 were also enriched in bacterial samples, 10 in viral samples, and 4 were unique to KD. Eight of the top 19 KD pathways were enriched in both bacterial and viral samples. All of these are involved in the immune response. The higher frequency of overlapping concordant pathways makes it harder to identify differences between the pathways enriched in the proteomic dataset than the transcriptomic dataset. Overall there was a much lower number of proteins SDA between KD vs HC (n = 113) than genes SDA between KD vs HC (n = 3,218). Furthermore, the total number of proteins remaining following quality control and filtering for missingness (n = 344) was much lower than the total number of genes remaining following quality control (n = 13,035), which could justify why the differences in pathways enriched between the disease groups are more apparent in the gene expression data.

Three pathways were enriched on both ‘omic levels. These were: immune effector process pathway (upregulated in KD and bacterial patients); immune response (upregulated in KD patients); and vesicle-mediated transport (upregulated in KD and bacterial patients).

#### 2.2.3. Clustering

*K*-Means clustering was used to determine whether the KD patients were more likely to cluster with bacterial or viral patients in the discovery datasets. Prior to clustering analysis, gene expression values were corrected for age, sex and immune cell proportions by taking the residual gene expression values after removing the contributions of these variables. Immune cell proportions were estimated using CIBERSORTx [25], an online tool for estimating immune cell proportions from gene expression data. The same process was performed to remove the contribution of age and sex from the protein abundance values. NbClust [26] was used to determine the optimal number of clusters (*k*). The value of *k* most frequently selected across the 12 indices measured by NbClust was selected as the optimal number of clusters for downstream analyses. In the transcriptomic analysis, 3 clusters were identified as optimal, whereas on the protein level, 2 clusters were identified as optimal.

We assessed the proportion of KD, bacterial and viral patients in each of the clusters for the transcriptomic (A) and proteomic (B) datasets (Fig. 2). In the transcriptomic analysis, an over-representation of viral patients was observed in cluster 1 and, to a lesser extent, cluster 2. An over representation of bacterial patients was observed in cluster 3 (Fig. 2), resulting in two viral-like clusters and one bacterial-like cluster. Of the 77 transcriptomic KD samples, 47 (61%) belonged to cluster 3, 22 (29%) belonged to cluster 2, and 8 (10%) belonged to cluster 1. In the proteomic analysis, an over-representation of bacterial patients was found in cluster 1, whereas an over-representation of viral patients was observed in cluster 2, leading to one viral-like and one bacterial-like cluster. Of the 26 proteomic KD samples, 24 (92%) belonged to cluster 1 and 2 (8%) belonged to cluster 2.

**Figure 2.**
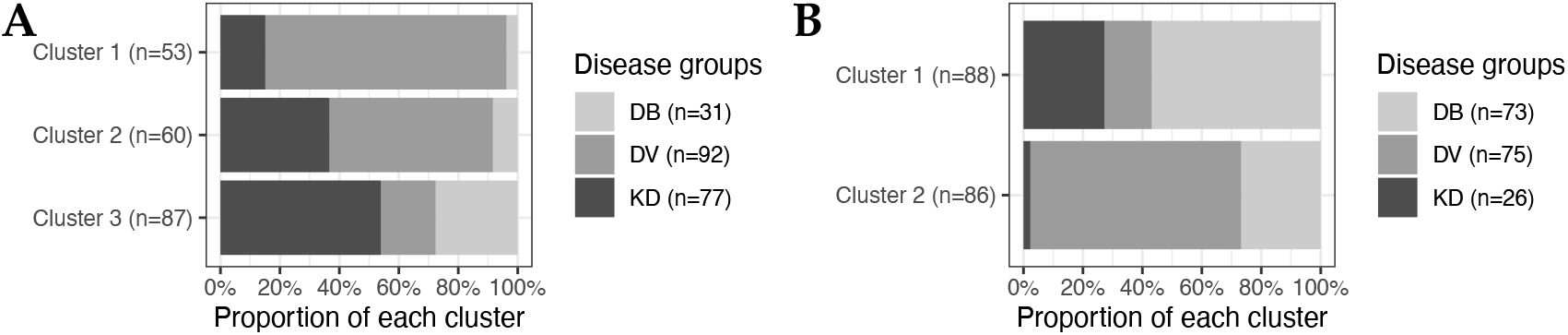
The proportion of patients from each disease group in each cluster for transcriptomics (A) and proteomics (B). DB, DV and KD represent definite bacterial, definite viral, and Kawasaki Disease.

The association between KD patient cluster membership and various clinical variables was tested. CRP levels (p-value: 0.048), lymph node swelling (p-value: 0.044), and peeling (p-value: 0.050) were significantly associated with cluster membership of KD transcriptomic samples. Higher levels of CRP were found in transcriptomic KD samples in cluster 3 which had the highest proportion of bacterial samples. Out of the 55 patients displaying peeling, 38 were found in cluster 3, as were 17 of the 21 patients with lymph node swelling. No clinical variables were associated with cluster membership in the proteomic dataset. On both ‘omic levels, CRP levels were highest in the clusters in which the majority of bacterial samples were found (Fig. S5-S6), as expected since a CRP cut-off of <60mg/L was required for patients in the DV groups. This pattern was also observed for the WBC counts in the transcriptomic dataset (Fig. S5).

Differential abundance analysis was performed to compare feature abundance in the KD samples that fell into different clusters. There were 503 genes SDA between transcriptomic KD samples in cluster 1 vs clusters 2 and 3, 454 genes SDA between KD samples in cluster 2 vs clusters 1 and 3, and 651 genes SDA between KD samples in cluster 3 vs clusters 1 and 2. These lists of SDA genes were subjected to pathway analysis using g:Profiler2 [24] to identify pathways upregulated and downregulated within the clusters (Fig. 3). Complete lists of pathways are in Supplementary File 3.

**Figure 3.**
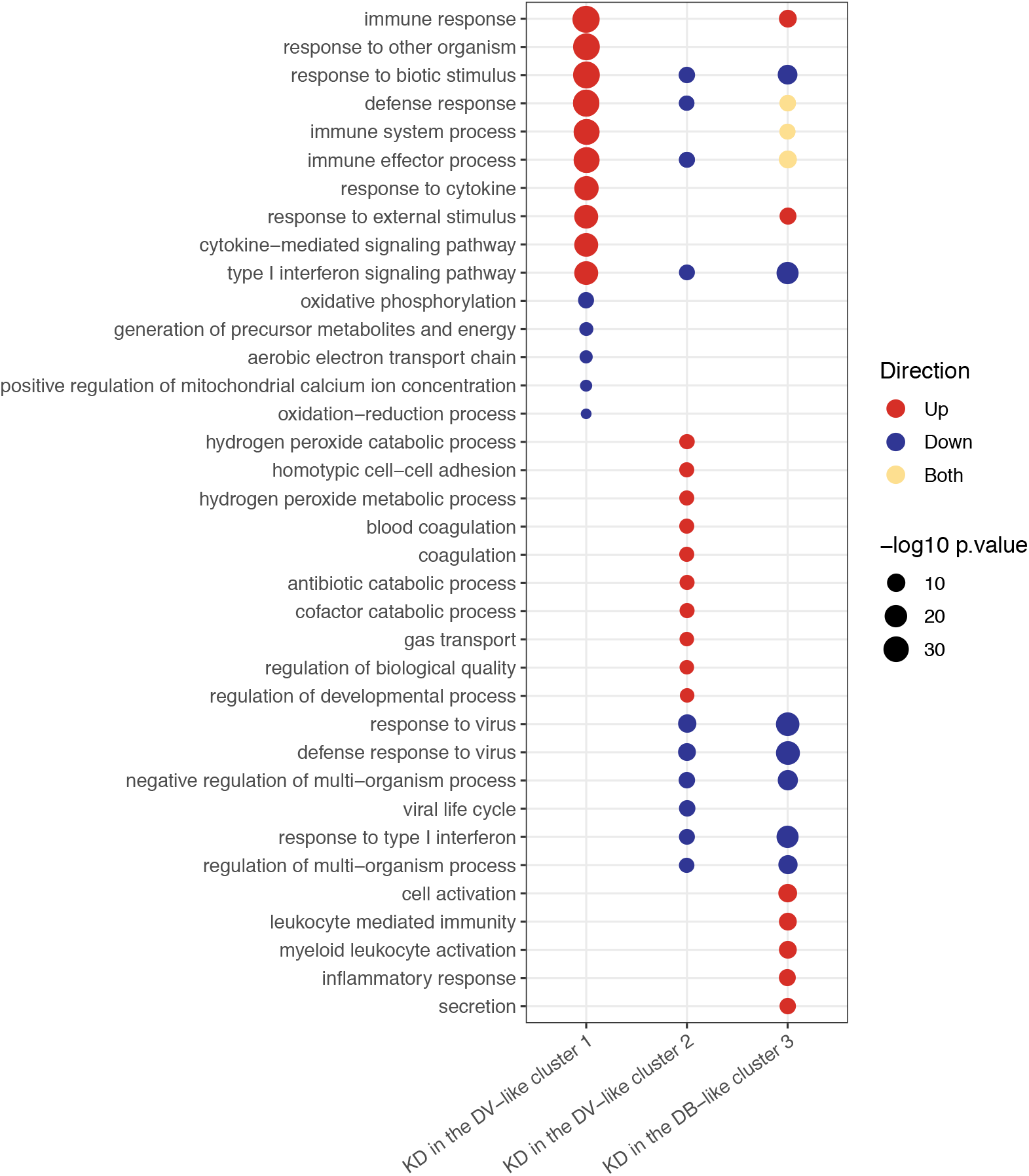
Pathways upregulated and downregulated in the KD patients in clusters 1, 2 and 3 for the transcriptomic dataset. Clusters were identified using *K*-Means applied to KD, DB and DV patients. There were 151, 52 and 137 pathways upregulated in clusters 1, 2 and 3, respectively, and 5, 66 and 137 pathways downregulated in clusters 1, 2 and 3, respectively.

For the transcriptomics, cluster 1 had the highest proportion of viral patients compared to the other clusters (Fig. 2a). The majority of the Adenovirus (19/23) and Influenza (16/23) patients were in cluster 1. Cluster 1 KD patients were characterised by upregulation of anti-viral response pathways, such as interferon and cytokine signalling (Fig. 3). In cluster 2, although the majority of patients were viral, their proportion was not quite as high as it was in cluster 1 (Fig. 2). The majority of the RSV (15/27) patients were in cluster 2. In the KD patients in cluster 2, various pathways associated with the anti-viral response were downregulated (Fig. 3). Cluster 3 had the highest proportion of bacterial patients and KD patients (Fig. 2). Similarly to cluster 2, the top pathways downregulated for KD patients in cluster 3 were associated with the anti-viral response, while the inflammatory response pathway was strongly upregulated suggesting that the KD patients in this cluster were different to those in cluster 1 and that their response was not as viral-like as those in cluster 1 (Fig. 3).

Three pathways - response to biotic stimulus (i.e. a stimulus caused or produced by a living organism), response to other organism and type I interferon signalling - were upregulated in viral transcriptomic samples (Fig. 1a) and also in the KD samples in the viral-like cluster 1 (Fig. 3). Furthermore, four pathways, including two associated with interferon signaling, were upregulated in viral transcriptomic samples (Fig. 1a) and downregulated in the KD samples in clusters 2 and 3 (Fig. 3). There were five pathways downregulated in bacterial transcriptomic samples (Fig. 1a) and upregulated in KD transcriptomic samples in cluster 1 (Fig. 3), including two related to cytokine signaling.

For the proteomic dataset, two proteins were SDA between cluster 1 and 2: serum amyloid A1 (SAA1) and retinol binding protein 4 (RBP4). Both of these proteins have been identified previously as Kawasaki markers, with RBP4 abundance being lower in active KD [27] and SAA1 being elevated in KD [28]. The two KD patients in cluster 2 displayed the opposite pattern, with higher RBP4 and lower SAA1 abundance than the other KD patients.

#### 2.2.4. Classification using Disease Risk Scores

To further assess if the KD patients elicited more bacterial-like or more viral-like responses, we built two classifiers that returned the probabilities that a patient is bacterial or viral through two separate disease risk scores (DRS). A DRS translates the abundance of features in a discriminatory signature, selected by Lasso [29], into a single value that can be assigned to each individual [16]. Through using two independent classifiers, the possibility of a patient being neither bacterial nor viral was allowed. The classifiers were trained using the ‘omic data that was corrected for age, sex and, for the transcriptomic dataset, immune cell proportions.

The Lasso model selected 38 genes for the bacterial classifier, of which 26 had increased abundance and 12 had decreased abundance in bacterial patients compared to viral patients and healthy controls (Table S4). The viral classifier included 32 genes, of which 13 had increased abundance and 19 had decreased abundance in viral patients compared to bacterial patients and healthy controls (Table S5). The classifiers trained in the transcriptomic discovery dataset were tested on bacterial and viral patients from the transcriptomic validation dataset. The bacterial classifier achieved an area under the ROC curve (AUC) of 0.935 (95% CI: 0.869-1) and the viral classifier achieved an AUC of 0.935 (95% CI: 0.856-1).

The Lasso model selected 26 proteins for the bacterial classifier, of which 12 had increased abundance and 14 had decreased abundance in bacterial patients compared to viral patients and healthy controls (Table S6). The viral classifier included 20 proteins, of which 11 had increased abundance and 9 had decreased abundance in viral patients compared to bacterial patients and healthy controls (Table S7). When testing the classifiers trained in the proteomic discovery dataset on bacterial and viral patients from the validation dataset, the bacterial classifier achieved an AUC of 0.925 (95% CI: 0.867-0.984) and the viral classifier achieved an AUC of 0.891 (95% CI: 0.821-0.962). For both ‘omic levels, the 90% sensitivity of the classifiers in classifying these samples was used to determine the DRS threshold above which a sample would be classified as bacterial or viral.

The classifiers were applied to KD patients from the discovery and validation datasets for both ‘omic levels, resulting in bacterial DRS (DB-DRS) and viral DRS (DV-DRS) for each KD patient (Fig. 4-5). Classification labels (DB or DV) were assigned to the KD patients using the DB-DRS and DV-DRS thresholds calculated from applying the classifiers to the bacterial and viral patients in the validation datasets (Fig. S9). Of the 178 transcriptomic KD samples, 18 (10%) samples had DB-DRS high enough to be classified as bacterial and 16 (9%) samples had DV-DRS high enough to be classified as viral. 145 (81%) samples did not achieve DB-DRS nor DV-DRS sufficiently high to lead to bacterial or viral classification, and 1 sample was classified as both bacterial and viral (Fig. 4). Of the 52 proteomic KD samples, 40 (78%) achieved DB-DRS high enough to be classified as bacterial and 18 (35%) achieved DV-DRS high enough to be classified as viral. 10 (19%) proteomic KD samples achieved DB-DRS and DV-DRS high enough for them to be classified as both bacterial and viral, and 4 (7.7%) were classified as neither (Fig. 5).

**Figure 4.**
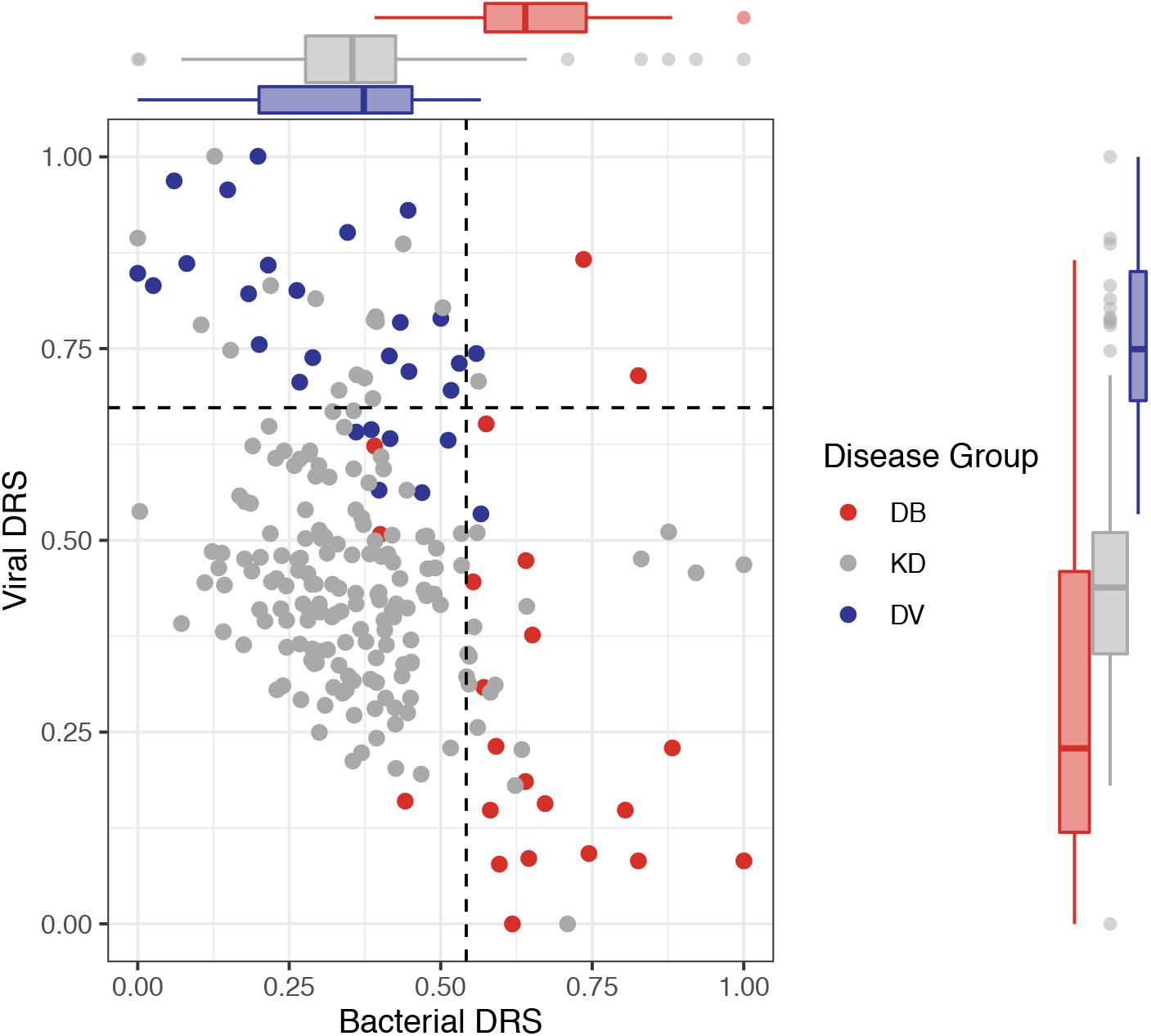
Bacterial DRS (DB-DRS) plotted against viral DRS (DV-DRS) for KD (discovery and validation), definite bacterial (DB; validation) and definite viral (DV; validation) patients from the transcriptomic datasets. Boxplots are shown for each disease group.

**Figure 5.**
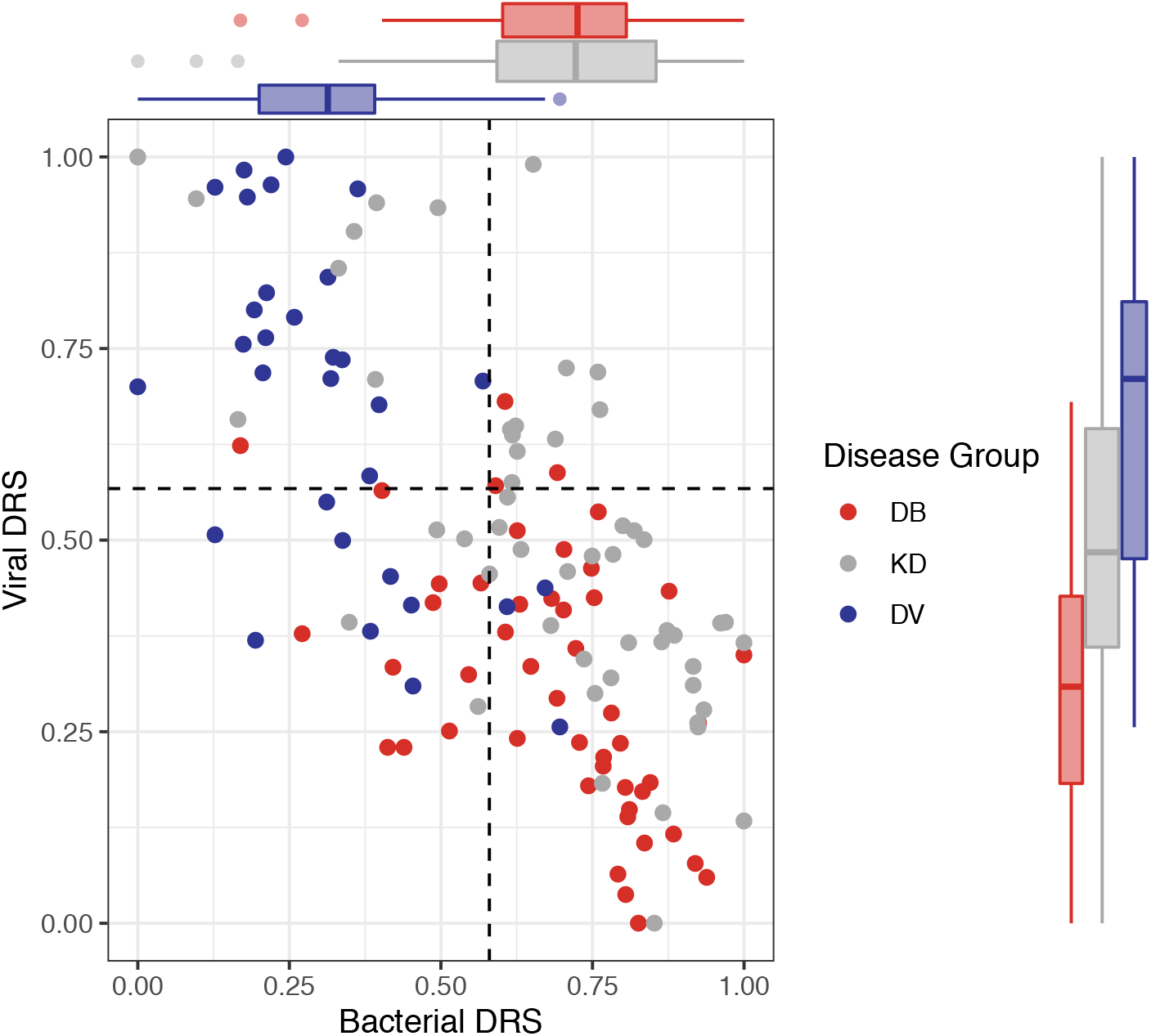
Bacterial DRS (DB-DRS) plotted against viral DRS (DV-DRS) for KD (discovery and validation), definite bacterial (DB; validation) and definite viral (DV; validation) patients from the proteomic datasets. Boxplots are shown for each disease group.

To further examine the ‘omic profiles of the KD patients with DB-DRS and DV-DRS too low for them to be classified as either bacterial or viral, we performed pathway analysis on the genes or proteins SDA between these KD patients and healthy controls. Amongst the pathways upregulated on the transcriptomic level, were ‘defense response to fungus’ (p-value: 6.6e-08) and and ‘response to fungus’ (p-value: 6.7e-07).

The associations of DB-DRS, DV-DRS, bacterial classification as predicted from the DB-DRS, and viral classification as predicted from the DV-DRS, with various clinical variables were tested for KD samples from both ‘omic levels. In the transcriptomic KD samples, clinical measurements of CRP were positively associated with DB-DRS (p-value: 0.002) and bacterial classification (p-value: 0.0001), and negatively associated with DV-DRS (p-value: 0.002) and viral classification (p-value: 0.023). In the proteomic KD samples, CRP levels were significantly positively associated with DB-DRS (p-value: 0.013) and bacterial classification (p-value: 0.007). Peeling was significantly associated with higher DB-DRS on both ‘omic levels (transcriptomic p-value: 0.041, proteomic p-value: 0.007). Strawberry tongue was significantly associated with a low score on the transcriptomic DV-DRS (p-value: 0.045).

For the KD patients from the discovery datasets, the associations between DB-DRS or DV-DRS and the cluster membership of patients was tested. Transcriptomic KD sample cluster membership was significantly associated with DB-DRS (p-value: 0.005) and DV-DRS (p-value: 0.0006), with a stepwise increase in DB-DRS and decrease in DV-DRS from clusters 1 to 3, where cluster 1 was the most viral-like cluster, and cluster 3 was the most bacterial-like cluster. Proteomic KD sample cluster membership was significantly associated with DB-DRS (p-value: 0.002) and DV-DRS (p-value: 0.023), with higher DB-DRS and lower DV-DRS in KD patients in cluster 1, where cluster 1 was the more bacterial-like cluster and cluster 2 was the more viral-like cluster.

### 2.3. Clustering of Kawasaki Disease patients alone

We performed unsupervised clustering for the KD patients from the discovery datasets to explore the natural patient stratification formed in the absence of bacterial and viral comparator patients. For both ‘omic levels, 3 clusters were optimal, as determined by NbClust [26]. The clusters were identified using the ‘omic data that was corrected for age, sex and, for the transcriptomic dataset, immune cell proportions. Of the 77 transcriptomic KD samples, 32 (41%) were in cluster 1 (cluster KD1-T), 23 (30%) were in cluster 2 (cluster KD2-T), and 22 (29%) were in cluster 3 (cluster KD3-T). Of the 26 proteomic KD samples, 4 (15%) were in cluster 1 (cluster KD1-P), 7 (27%) were in cluster 2 (cluster KD2-P), and 15 (58%) were in cluster 3 (cluster KD3-P).

There was high overlap between the samples in cluster KD1-T and those in the transcriptomic bacterial-like cluster 3 described in *2.2.3* (Fig. S10). All except one of the samples found previously in the transcriptomic viral-like cluster 1 were found in cluster KD2-T. The majority (n = 14; 64%) of the samples in KD3-T were also found in transcriptomic cluster 2. On the proteome level, in *2.2.3*, all KD samples except two clustered together in cluster 1, however the two remaining samples that were previously in cluster 2, were not assigned to the same cluster.

The association between cluster membership and various clinical variables was tested. CRP levels were significantly associated with cluster membership for both ‘omic layers (transcriptomics p-value: 0.041, proteomics p-value: 0.010). Furthermore, coronary artery aneurysm (CAA) formation was significantly associated with cluster membership in the proteomic dataset (p-value: 0.020) with 13 of the 21 patients known to not have CAAs being in cluster KD3-P. On the transcriptomic level, the highest WBC counts and CRP levels were in cluster KD1-T, and on the proteomic level, WBC counts and CRP levels were highest in clusters KD2-P and KD1-P, respectively (Fig. S7-S8).

The associations between DB-DRS or DV-DRS and cluster membership of KD patients when clustered alone was tested. The transcriptomic KD samples’ cluster membership was significantly associated with DB-DRS (p-value: 0.006) with the highest DB-DRS in cluster KD1-T. Although the association between transcriptomic KD samples’ cluster membership and DV-DRS was not significant, the highest DV-DRS values were observed in cluster KD2-T. There were no signficant associations between the proteomic KD samples’ cluster membership and DB-DRS or DV-DRS.

Differential abundance analysis was performed on the patients that fell into different clusters. For the transcriptomics, there were 494 genes SDA between cluster KD1-T vs clusters KD2-T and KD3-T, 461 genes SDA between cluster KD2-T vs clusters KD1-T and KD3-T, and 320 genes SDA between cluster KD3-T vs clusters KD1-T and KD2-T. For the proteomics, 42 proteins were SDA between cluster KD1-P vs clusters KD2-P and KD3-P, 25 proteins were SDA between cluster KD2-P vs clusters KD1-P and KD3-P, and 38 proteins were SDA between cluster KD3-P vs clusters KD1-P and KD2-P. These lists of SDA features were subjected to pathway analysis using g:Profiler2 [24] to identify pathways upregulated and downregulated within the clusters (Fig. 6). Complete lists of pathways are found in Supplementary Files 4-5.

**Figure 6a.**
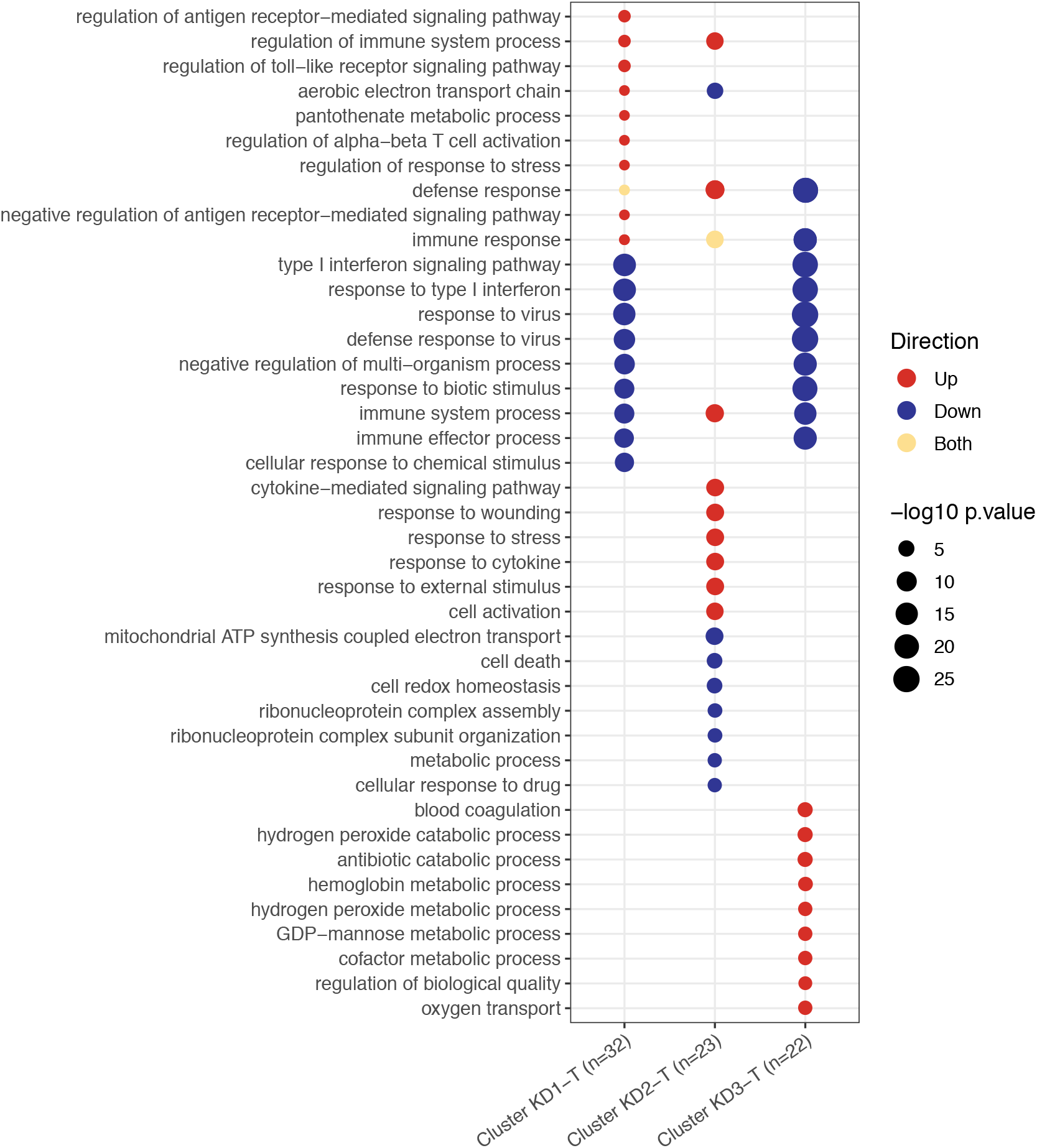
Pathways upregulated and downregulated in transcriptomic KD patients between clusters. Clusters were identified by *K*-Means ran on KD patients alone. There were 24, 118 and 24 pathways upregulated in clusters KD1-T, KD2-T and KD3-T, respectively, and 94, 68 and 75 pathways downregulated in clusters KD1-T, KD2-T and KD3-T, respectively.

**Figure 6b.**
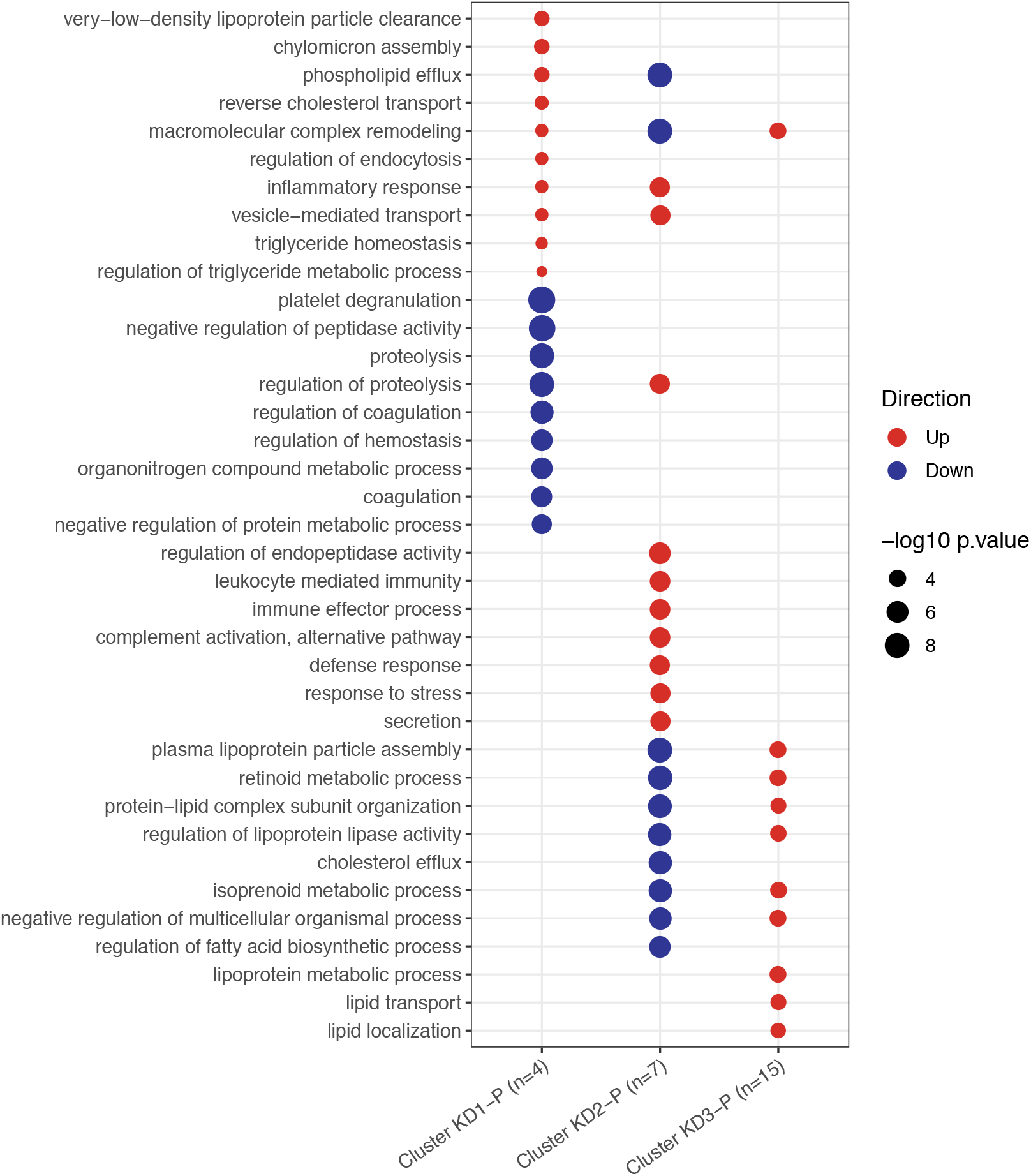
Pathways upregulated and downregulated in proteomic KD patients between clusters. Clusters were identified by *K*-Means ran on KD patients alone. There were 77, 94 and 61 pathways upregulated in clusters KD1-P, KD2-P and KD3-P, respectively, and 64, 104 and 53 pathways downregulated in clusters KD1-P, KD2-P and KD3-P, respectively.

In the transcriptomic analysis (Fig. 6a), cluster KD2-T had features in common with an anti-viral response, whilst the others did not. Many pathways associated with the anti-viral response were downregulated in clusters KD1-T and KD3-T whilst patients in cluster KD2-T were characterised by the upregulation of viral pathways, including those associated with cytokine signaling.

The response to biotic stimulus and type I interferon signaling pathways were previously identified as being upregulated in viral transcriptomic samples (Fig. 1a) and in KD samples in the viral-like cluster 1 when *K-*Means was applied to KD, DB and DV (Fig. 3). These pathways were downregulated in clusters KD1-T and KD3-T (Fig. 6a), indicating that the transcriptomic response in these samples was less viral-like than samples in cluster KD2-T.

Four pathways previously identified as being downregulated in bacterial transcriptomic samples (Fig. 1a), including two pathways associated with cytokine signaling, were upregulated in cluster KD2-T. In addition, six pathways upregulated in cluster KD2-T had already been identified as being upregulated in the viral-like cluster 1 identified previously (Fig. 3). Five pathways of the 9 top upregulated pathways in cluster KD3-T (Fig. 6a) were also upregulated in cluster 2 when *K*-Means was applied to KD, DB and DV (Fig. 3). These were blood coagulation, hydrogen peroxide catabolic process, antibiotic catabolic processes, hydrogen peroxide metabolic process and regulation of biological quality.

In the proteomic analysis (Fig. 6b), one of the KD clusters had features in common with the anti-viral response, whilst another KD cluster was more bacterial-like. Amongst the top pathways enriched in cluster KD1-P and KD2-P were pathways involved in inflammation. The top pathways enriched in cluster KD3-P were associated with lipids. Of the 37 pathways enriched in the proteomic KD samples (Fig. 6b), 21 were previously identified as enriched in proteomic samples (Fig. 1b). Of these, 6 were enriched in proteomic viral samples (Fig. 1b) and samples in cluster KD2-P (Fig. 6b) with concordant directions. Furthermore, 7 pathways enriched in cluster KD1-P (Fig. 6b) were also enriched in bacterial proteomic samples (Fig. 1b) with concordant directions. These results suggest that cluster KD1-P is a more bacterial-like cluster, whereas cluster KD2-P is a more viral-like cluster. Some pathways were enriched on both ‘omic levels, including those associated with blood coagulation, the response to stress and immune effector processes.

## 3. Discussion

Although the cause of Kawasaki Disease has not been identified, there is growing clinical, epidemiological and immunological evidence that it may be caused by different infectious triggers, with data pointing to bacteria, viruses or fungi. We explored the transcriptomes and proteomes of children with KD and definite bacterial and viral infections, using multiple approaches to compare the host response to these diseases to the response during KD. We found that there was a diversity of responses in the proteomic and transcriptomic profiles of KD patients, suggesting that KD is not a homogenous condition, and that whilst some patients had a more viral- or bacterial-like profile, the majority were defined as neither bacterial nor viral when their transcriptomic response was mapped onto viral and bacterial disease risk scores (DRS).

Within the host response profiles, some elements of KD appeared more viral-like and some elements appeared more bacterial-like. This is shown through overlapping pathways that were enriched in KD and either bacterial or viral infections. For example, the antigen presentation via MHC class I pathway was upregulated in KD and viral infections on the transcriptomic level. Major histocompatibility complex (MHC) molecules are expressed on the cell surface to present antigenic peptides to T cells, and their expression is increased by a broad range of immune activators including interferons [30,31]. The finding of upregulated MHC class I expression in KD and viral patients may reflect interferon-induced activation in these groups. Also, on the transcriptomic level, KD and bacterial infections share neutrophil degranulation as their most upregulated pathway. Neutrophils are the first responders to infection and inflammation, and the expansion and activation of the neutrophil population is a characteristic feature of acute KD. In the initial days of KD illness there is an intense inflammatory response with neutrophil leucocytosis [32]. Studies have found elevated levels of human neutrophil elastase and IL-8, a C-X-C chemokine that activates neutrophils [33,34].

The host response during KD is highly heterogenous, as demonstrated through the enrichment of certain pathways in the KD patients in different clusters when *K*-Means was applied to KD, bacterial and viral transcriptomic samples. For example, anti-viral response pathways were upregulated in KD patients in the majority viral cluster 1 and downregulated in KD patients in clusters with decreasing numbers of viral samples (cluster 2, 3), relative to each other. In the majority bacterial cluster 3, pathways associated with the inflammatory response were upregulated. The heterogeneity of the host response during KD was also apparent when *K*-Means was applied to KD patients alone. Three distinct clusters were identified on both ‘omic levels, and, in each cluster, a distinct set of pathways was enriched. The range of pathways enriched in the different clusters further demonstrate the heterogeneity in the host response during KD, with some clusters enriched for viral response pathways and some clusters enriched for bacterial response pathways. Unsurprisingly, amongst the patients clustering in the more bacterial-like clusters, their DB-DRS tended to be higher, and amongst the patients clustering in the more viral-like clusters, their DV-DRS tended to be higher.

The two different approaches to clustering (with and without bacterial and viral comparator samples) produced similar clusters of KD patients, providing reassurance that the clusters described here are biologically meaningful. Despite the similarities, however, the clusters identified in *2.2.3* and *2.3* were not completely identical, indicating that the inclusion of well characterized bacterial and viral patients adds further insights to the solely data-driven KD-based analysis.

Although there are shared features between the response to KD and both bacterial and viral infections, the distinct pathways enriched in each disease group demonstrate the variation in the molecular host response; the distinctiveness of the responses is also supported by the ability of RNA and protein signatures to discriminate KD from bacterial and viral infections [18,19,35]. These differences between the response during KD and the responses to bacterial and viral infections suggest that KD may be triggered by a novel process not typical of either common bacterial or viral infections. Despite the host omics profiles’ heterogeneity observed in KD, commonalities are also shown.

A two-way classifier approach highlighted that it is not a simple dichotomous question as to whether the response during KD more closely resembles the responses to bacterial or viral infections, when focusing on key discriminatory features. We found that 145 of the 178 transcriptomic KD samples were not assigned DRS high enough for them to be classified as either bacterial or viral, and amongst pathways upregulated in these KD patients compared to healthy controls were two pathways associated with the fungal response. This finding is intriguing, given the evidence suggesting that KD could be caused by a fungal trigger that has been reported elsewhere [36,37].

The heterogeneity and the different clusters of responses to KD which have elements shared with bacterial, viral or fungal responses, could indicate multiple microbial triggers of KD, as has been suggested by Rypdal et al. [11]. An alternative explanation for the heterogeneity observed here in the response during KD could be that a single pathogen that causes KD leads to heterogeneous responses in different hosts, as has been observed in children infected with SARS-CoV-2, where many children remain asymptomatic, some experience severe inflammation [38,39], and some develop PIMS-TS/MIS-C [13,14,40]. Variations in the host condition, such as epigenetic differences and differences in prior pathogen exposure, could cause the spectrum of host responses to KD observed here. Differences in host genetics could also be responsible for the heterogeneity in host response during KD as the severity of KD, including the formation of CAAs, is already known to be impacted by the host’s genetic background [41].

This study has certain limitations. The proteomic discovery dataset was a lower dimensional dataset (n= 867) than the transcriptomic discovery dataset (n=47,323) with high rates of missingness, as is common in quantitative proteomics. Only proteins with no missingness were used for the clustering and classification, so key proteins for distinguishing KD could be absent from the analysis. On the proteomic level, many pathways were enriched in multiple disease groups (Fig. 1b), making it difficult to identify a disease-specific pathway signature. This could be caused by plasma samples, which were used in this dataset, capturing a noisy signal due to the release of substances from various tissues into the bloodstream. The proteomic response during KD shared more similarities with the proteomic response to bacterial infection, with more pathways overlapping between KD and bacterial infections (Fig. 1b) and all but two KD proteomic samples clustering with bacterial proteomic samples (Fig. 2). This follows observations of striking clinical similarities between KD and bacterial streptococcal and staphylococcal toxic shock syndromes [42,43], and could reflect the hypothesis that the proteome is closer to the observed phenotype than the transcriptome [44].

Although bacterial patients with known viral coinfections have been removed from the analysis, it is impossible to say with confidence that an individual does not have a coinfection, the presence of which could falsely increase heterogeneity in the host response in a given disease group. Coinfection is common in KD, with one study identifying confirmed infections in a third of KD patients [45]. Despite being unable to rule-out that some KD patients included had co-incident viral or bacterial infections, we found that most KD transcriptomic samples were neither classified as viral nor bacterial when the respective DRS scores were applied. Amongst the patients classified as bacterial or viral, it is possible that some patients could be suffering from an intercurrent infection in addition to KD.

There are variations in the range of bacterial and viral pathogens and the severity of illness represented in the two ‘omic datasets. The bacterial and viral patients included in the transcriptomic datasets and the proteomic validation dataset were more severely unwell than those included in the proteomic discovery dataset (Table S3) due to the inclusion criteria of the studies to which they were recruited. The KD patients included in the transcriptomic dataset were collected from San Diego, USA, whereas the KD patients included in the proteomic dataset were collected from London, UK, although the same case definition was used. There remains no diagnostic test for KD, thus some KD patients presented here may have unrecognized alternative diagnoses. The 2 KD samples in the proteomic dataset that cluster separately (Fig. 2b) and are distinguished from the other KD samples by their levels of SAA1 and RBP4, two previously identified KD markers [27,28], are a possible example of this.

## 4. Materials and Methods

### 4.1. Patient recruitment

All samples were obtained from patients with written parental informed consent. Case definitions can be found in the Supplementary Text. The definite bacterial (DB), definite viral (DV), healthy control (HC) and Kawasaki Disease (KD) samples used in the transcriptomic discovery and validation datasets were recruited in the United Kingdom and Spain as part of the IRIS (Immunopathology of Respiratory, Inflammatory and Infectious Disease; NIHR ID 8209) and GENDRES (Genetic, Vitamin D, and Respiratory Infections Research Network; http://www.gendres.org) studies [17,22] and in the United States through the US-Based Kawasaki Disease Research Center Program (https://medschool.ucsd.edu/som/pediatrics/research/centers/kawasaki-disease/pages/default.aspx).

The DB, DV and HC samples used in the proteomic discovery and validation datasets were enrolled in the EUCLIDS (European Union Childhood Life-Threatening Infectious Disease Study; 11/LO/1982) study [46] and the PERFORM (Personalised Risk assessment in Febrile illness to Optimise Real-life Management across the European Union) study (https://www.perform2020.org/; 16/LO/1684). KD samples used in the proteomic datasets were recruited from the ongoing UK Kawasaki study “Genetic determinants of Kawasaki Disease for susceptibility and outcome” (13/LO/0026). This study recruits acutely unwell children with KD during hospital admission in participating hospitals around the UK.

### 4.2. Data generation

#### 4.2.1. Transcriptomic datasets

The transcriptomic discovery dataset was generated from whole blood samples obtained from KD patients, healthy controls, and patients with bacterial and viral infections using the HumanHT-12 version 4.0 (Illumina) microarray[18]. In order to have a transcriptomic validation dataset containing the same disease groups as the transcriptomic discovery dataset, two datasets were merged. One dataset contained gene expression values (HumanHT-12 version 4.0 Illumina microarray) from whole blood samples obtained from acute and convalescent KD samples [19]. The other dataset consisted of gene expression values (HumanHT-12 version 3.0 Illumina microarray) from whole blood samples obtained from patients with bacterial and viral infections [22]. For all three independent microarray experiments, one batch of samples was processed, and samples were randomly positioned across the arrays.

#### 4.2.1. Proteomic datasets

The proteomic discovery dataset was generated from plasma samples using LC-MS/MS. Full details of the experimental protocol are in the Supplementary Text. The proteomic validation dataset was generated from serum samples using the SomaScan aptamer-based platform [20]. Prior to pre-processing, 867 proteins were measured in the discovery dataset (LC-MS/MS) and 1,300 in the validation dataset (SomaScan). Samples in the proteomic validation dataset were split across three plates with KD, DB, DV and HC samples present on each plate in relative proportions.

### 4.3. Statistical methods

All analysis was conducted using the statistical software R (R version 3.6.1, [49]). Code used for the analytical pipeline described here is found at https://github.com/heather-jackson/KawasakiDisease_IJMS. Note, the code is signposted for the transcriptomic datasets but can be modified for other ‘omic levels.

#### 4.3.1. Pre-processing of gene expression data

Background correction, robust spline normalisation (RSN), and log2-transformation were applied to the raw discovery gene expression dataset using the R package lumi [47]. Probes were retained if at least 80% of samples in each comparator group had a detection p-value <0.01. Low variance probes and those significantly associated with the UCSD recruitment site were removed. Bacterial samples with known viral coinfections were removed from the analysis at this stage to ensure that the signal from the bacterial samples was not diluted. KD samples that had been administered IVIG treatment were also removed at this stage, but their inclusion was irrespective of coincident viral or bacterial detection, for which data was not available. A KD sample previously identified as an outlier [18] was removed.

As mentioned, two microarray gene-expression datasets were merged to form the validation dataset. Background subtraction and RSN normalisation were applied to these two datasets independently, using the R package lumi [47], prior to using ComBat [48] to remove the batch effects in the merged dataset [18].

#### 4.3.2. Pre-processing of protein abundance data

The raw discovery dataset files generated by LC-MS/MS were processed using MaxQuant (1.6.10.43) [50]. with matching between runs activated. Relative quantification was performed using the MaxLFQ algorithm [51]. The resulting LFQ values were log2-transformed. Bacterial samples with known viral coinfections were removed from the analysis at this stage to ensure that the signal from the bacterial samples was not diluted. Protein groups were removed if they were identified as contaminants, or if they were missing in over 90% of samples in each disease group.

The proteomic validation dataset was generated from the SomaScan platform [20]. Quality control steps used scale factors returned from SomaScan to correct for variations in aptamer hybridisation efficiency, inter- and intra-assay variability, variability in the starting quantities of proteins, and plate effects. Further batch effect corrections were carried out using COCONUT normalisation [52].

#### 4.3.3. Comparison of Kawasaki Disease to bacterial and viral infections

##### 4.3.3.1. Differential abundance analysis

Differential abundance analysis was carried out to compare the overall transcriptomic and proteomic responses to KD, definite bacterial (DB) and definite viral (DV) infections. The degree to which genes and proteins were differentially abundant between KD and healthy controls (HC), DB and HC, and DV and HC was quantified using Limma [23] on the transcriptomic and proteomic discovery datasets separately. Age and sex were included as covariates for both datasets. Immune cell proportions, calculated using the online CIBERSORTx portal [25], were used as additional covariates for the transcriptomic dataset. The immune cell proportions included were lymphocytes, neutrophils, monocytes, mast cells and eosinophils. Features were considered significantly differentially abundant (SDA) at a false discovery rate (FDR) [53] of 5%.

##### 4.3.3.2. Pathway analysis

The pathways upregulated and downregulated in KD, DB, and DV samples were identified from the lists of SDA features identified for each disease group in as outlined in *4.3.3.1*. Pathways were identified using g:Profiler2 [24] and redundancy in the pathways identified was removed using REVIGO [54].

##### 4.3.3.3. Clustering analysis

*K-*Means clustering [55] was applied separately to transcriptomic and proteomic discovery datasets. Healthy controls were excluded as we were only interested in the clustering of KD with pathological patients. For the proteomic dataset, only proteins with no missing data points were used (n = 106).

To explore the effects of sex and age on clustering in the proteomic dataset, the contribution of these variables was removed by regressing out their effects on every protein and taking the residual values as the ‘corrected’ abundance. This process was also followed in the transcriptomic dataset but the contributions of the immune cell proportions listed in *4.3.3.1* were also removed. Prior to clustering, and after correction, features were removed if their variance was lower than 0.25. To determine the optimal number of clusters (*k*) for each corrected and non-corrected dataset, the R package NbClust [26] was used, with 12 indicies tested. The indices tested were: KL [56], CH [57], Hartigan [58], McClain [59], Dunn [60], SDIndex [61], SDbw [62], C-Index [63], Silhouette [64], Ball [65], Ptbiserial [66,67] and Ratkowsky [68]. The number of clusters tested by NbClust ranged between 2-10 clusters. The most frequently selected *k* by the 12 indices was used for downstream analyses. The lowest *k* selected the most frequently was taken in cases where there were multiple values of *k* selected the most frequently.

Once clusters were identified, features that were SDA (5% FDR) between KD samples in the different clusters were identified. Pathway analysis was done using these lists of SDA features to determine the pathways upregulated and downregulated in KD samples in the different clusters. The R package g:Profiler2 [24] was used for pathway analysis, with pathways with p-values < 0.01 considered significant. Redundancy in the pathways identified was removed using REVIGO [54].

The association between cluster membership and various clinical variables was tested. For categorical variables, Fisher’s Exact test was used. For continuous variables, One-Way ANOVA was used. P-values < 0.05 were considered significant. The categorical variables tested in both datasets were: strawberry tongue (yes/no/unknown); lymph node swelling (yes/no/unknown); and peeling (yes/no). Continuous variables tested in both datasets were: levels of C-reactive protein (CRP); month of year; and the duration of fever at sampling. Coronary artery aneurysms (CAA) was available only as a dichotomous variable for the patients submitted for proteomic analysis. For the patients submitted for transcriptomic analysis, maximal coronary artery Z-scores were available and were used instead.

##### 4.3.3.4. Classification

Two independent classifiers were built for each ‘omic dataset. One classifier was for classifying DB patients (DB classifier), and the other for classifying DV patients (DV classifier). The DB classifiers and DV classifiers were trained on features SDA between DB vs DV and HC patients combined, and DV vs DB and HC patients combined, respectively. Only features present in both datasets (discovery and validation) were used for training the classifiers. The discovery datasets that were corrected for age, sex, and, for transcriptomics, immune cell proportions were used to train the classifiers. The validation datasets were also corrected for age, sex and, for the transcriptomic validation dataset, immune cell proportions as determined by CIBERSORTx [25]. The proteomic discovery and validation datasets were generated using different platforms. Therefore, each dataset was scaled so that all abundance values were between 0-1, and then the two datasets were quantile normalised together. Proteins with no missing values that were also found in the proteomic validation dataset were used to train the proteomic classifier.

The DB classifiers were trained to identify DB patients from DV and HC patients, whereas the DV classifiers were trained to identify DV patients from DB and HC patients. Lasso regularised regression [29] was used to identify the discriminatory features and their weights for each classifier. For each sample, a disease risk score (DRS) was calculated using the abundance of the features selected by Lasso, as described by Kaforou et al. in [16]. The DRS was calculated by totalling the abundance of features with positive Lasso weights and subtracting from this total the abundance of features with negative Lasso weights. Features were only included in the DRS if their Lasso weight direction and log-fold change direction were concordant. DRS were scaled between 0-1.

The classifiers were tested on the DB and DV patients of their respective validation dataset. The cut-off threshold above which a sample was classified as DB or DV was calculated using the coords function in the R package pROC [69] using a sensitivity cut-off of 90%. The classifiers were then tested on the KD patients from the discovery and validation datasets and the thresholds identified by pROC were used to determine if the KD patients were classified as DB or DV. If patients were classified as neither bacterial nor viral according to their DRS, differential abundance analysis followed by pathway analysis (as described in *4.3.3.2*) was done to identify the pathways enriched in these patients compared to healthy controls.

#### 4.3.4. Exploration of Kawasaki Disease samples alone

In order to identify the natural clusters formed by KD patients in the absence of bacterial or viral patients, *K*-Means clustering was done separately on the KD patients. The process followed was the same as outlined in *4.3.3.3*. The association between cluster membership and clinical variables was tested. The clinical variables and the statistical tests used were the same as outlined in *4.3.3.3*.

## 5. Conclusions

Taken together, the results from differential abundance analysis, pathway analysis, clustering and classification suggest that the host transcriptomic and proteomic responses during KD are highly heterogenous. Different clusters of host responses during KD were identified, some of which resemble elements of host responses to bacterial, viral and fungal infections. These differences in the host responses could imply that KD is triggered either by several different pathogens, or by a single pathogen that has different manifestations according to the underlying genetic and environmental situation of the host. Whilst there are similarities between the host response during KD and the host response to bacterial infections and viral infections, there are also many differences in the responses, suggesting that KD may be triggered by a novel process not typical of either common bacterial or viral infections. This was demonstrated by the majority of the KD transcriptomic samples falling into a non-bacterial, non-viral group following classification, raising the possibility that the minority of KD transcriptomic samples with bacterial or viral profiles were possibly suffering from intercurrent infection in addition to a separate KD trigger. Our data further suggest that research into the etiologies of KD should be focused on cohorts of KD patients who share similar clinical characteristics in order to identify shared molecular responses.

## Author Contributions

Conceptualization: MK and ML; methodology: SH, HJ, AM, SM, CCP, MK; formal analysis: HJ, AM, MK; investigation: SH, MJ, SM, CCP, CS, VW; resources: JCB, RG, SH, MJ, SM, CS, AHT, VW; data curation: JCB, SH, JH, HJ, AM, SM, CS, VW; writing—original draft preparation: HJ; writing—review and editing: ALL; validation: HJ; visualization: HJ; supervision: JH, ML, MK; project administration: HJ, CS, VW, MK; funding acquisition: HJ, SM, SH, RG, AHT, MJ, TK, VW, JCB, CP, JH, ML, MK. All authors have read and agreed to the published version of the manuscript.

## Acknowledgments

Many thanks to the patients and their families who took part in the studies that the data presented here originated from.

## Abbreviations

CAA: Coronary artery aneurysm
DB: Definite bacterial
DRS: Disease risk score
DV: Definite viral
FDR: False discovery rate
HC: Healthy control KD Kawasaki Disease
LFC: Log-fold change
SDA: Significantly differentially abundant

## References

1. Kawasaki, T., Kosaki, F., Okawa, S., Shigematsu, I., Yanagawa, H. A New Infantile Acute Febrile Mucocutaneous Lymph Node Syndrome (MLNS) Prevailing in Japan. Pediatrics 1974, 54.

2. Ramphul, K., Mejias, S.G. Kawasaki disease: a comprehensive review. Arch. Med. Sci. - Atheroscler. Dis. 2018, 3, 41–45, doi:10.5114/amsad.2018.74522.

3. Ogata, S., Shimizu, C., Franco, A., Touma, R., Kanegaye, J.T., Choudhury, B.P., Naidu, N.N., Kanda, Y., Hoang, L.T., Hibberd, M.L., et al. Treatment Response in Kawasaki Disease Is Associated with Sialylation Levels of Endogenous but Not Therapeutic Intravenous Immunoglobulin G. PLoS One 2013, 8, e81448, doi:10.1371/journal.pone.0081448.

4. Skochko, S.M., Jain, S., Sun, X., Sivilay, N., Kanegaye, J.T., Pancheri, J., Shimizu, C., Sheets, R., Tremoulet, A.H., Burns, J.C. Kawasaki Disease Outcomes and Response to Therapy in a Multiethnic Community: A 10-Year Experience. J. Pediatr. 2018, 203, 408-415.e3, doi:10.1016/j.jpeds.2018.07.090.

5. Brogan, P., Burns, J.C., Cornish, J., Diwakar, V., Eleftheriou, D., Gordon, J.B., Gray, H.H., Johnson, T.W., Levin, M., Malik, I., et al. Lifetime cardiovascular management of patients with previous Kawasaki disease. Heart 2020, 106, 411–420.

6. Singh, S., Vignesh, P., Burgner, D. The epidemiology of Kawasaki disease: A global update. Arch. Dis. Child. 2015, 100, 1084–1088.

7. Nagata, S. Causes of Kawasaki Disease—From Past to Present. Front. Pediatr. 2019, 7, 18, doi:10.3389/fped.2019.00018.

8. Dietz, S.M., van Stijn, D., Burgner, D., Levin, M., Kuipers, I.M., Hutten, B.A., Kuijpers, T.W. Dissecting Kawasaki disease: a state-of-the-art review. Eur. J. Pediatr. 2017, 176, 995–1009.

9. Nakamura, A., Ikeda, K., Hamaoka, K. Aetiological significance of infectious stimuli in Kawasaki disease. Front. Pediatr. 2019, 7, 244.

10. Rodó, X., Ballester, J., Cayan, D., Melish, M.E., Nakamura, Y., Uehara, R., Burns, J.C. Association of Kawasaki disease with tropospheric wind patterns. Sci. Rep. 2011, 1, pdoi:10.1038/srep00152.

11. Rypdal, M., Rypdal, V., Burney, J.A., Cayan, D., Bainto, E., Skochko, S., Tremoulet, A.H., Creamean, J., Shimizu, C., Kim, J., et al. Clustering and climate associations of Kawasaki Disease in San Diego County suggest environmental triggers. Sci. Rep. 2018, 8, 1–9, doi:10.1038/s41598-018-33124-4.

12. Levin, M. Childhood Multisystem Inflammatory Syndrome — A New Challenge in the Pandemic. N. Engl. J. Med. 2020, 383, 393–395, doi:10.1056/NEJMe2023158.

13. Whittaker, E., Bamford, A., Kenny, J., Kaforou, M., Jones, C.E., Shah, P., Ramnarayan, P., Fraisse, A., Miller, O., Davies, P., et al. Clinical Characteristics of 58 Children with a Pediatric Inflammatory Multisystem Syndrome Temporally Associated with SARS-CoV-2. JAMA - J. Am. Med. Assoc. 2020, 324, 259–269, doi:10.1001/jama.2020.10369.

14. Dufort, E.M., Koumans, E.H., Chow, E.J., Rosenthal, E.M., Muse, A., Rowlands, J., Barranco, M.A., Maxted, A.M., Rosenberg, E.S., Easton, D., et al. Multisystem Inflammatory Syndrome in Children in New York State. N. Engl. J. Med. 2020, 383, 347–358, doi:10.1056/NEJMoa2021756.

15. McCrindle, B.W., Manlhiot, C. SARS-CoV-2-Related Inflammatory Multisystem Syndrome in Children: Different or Shared Etiology and Pathophysiology as Kawasaki Disease? JAMA - J. Am. Med. Assoc. 2020, 324, 246–248.

16. Kaforou, M., Wright, V.J., Oni, T., French, N., Anderson, S.T., Bangani, N., Banwell, C.M., Brent, A.J., Crampin, A.C., Dockrell, H.M., et al. Detection of Tuberculosis in HIV-Infected and -Uninfected African Adults Using Whole Blood RNA Expression Signatures: A Case-Control Study. PLoS Med. 2013, 10, e1001538, doi:10.1371/journal.pmed.1001538.

17. Kaforou, M., Herberg, J.A., Wright, V.J., Coin, L.J.M., Levin, M. Diagnosis of Bacterial Infection Using a 2-Transcript Host RNA Signature in Febrile Infants 60 Days or Younger. JAMA 2017, 317, pdoi:10.1001/jama.2017.1365.

18. Wright, V.J., Herberg, J.A., Kaforou, M., Shimizu, C., Eleftherohorinou, H., Shailes, H., Barendregt, A.M., Menikou, S., Gormley, S., Berk, M., et al. Diagnosis of Kawasaki Disease Using a Minimal Whole-Blood Gene Expression Signature. JAMA Pediatr. 2018, 172, e182293, doi:10.1001/jamapediatrics.2018.2293.

19. Hoang, L.T., Shimizu, C., Ling, L., Naim, A.N.M., Khor, C.C., Tremoulet, A.H., Wright, V., Levin, M., Hibberd, M.L., Burns, J.C. Global gene expression profiling identifies new therapeutic targets in acute Kawasaki disease. Genome Med. 2014, doi:10.1186/s13073-014-0102-6.

20. Gold, L., Ayers, D., Bertino, J., Bock, C., Bock, A., Brody, E.N., Carter, J., Dalby, A.B., Eaton, B.E., Fitzwater, T., et al. Aptamer-based multiplexed proteomic technology for biomarker discovery. PLoS One 2010, doi:10.1371/journal.pone.0015004.

21. McCrindle, B.W., Rowley, A.H., Newburger, J.W., Burns, J.C., Bolger, A.F., Gewitz, M., Baker, A.L., Jackson, M.A., Takahashi, M., Shah, P.B., et al. Diagnosis, treatment, and long-term management of Kawasaki disease: A scientific statement for health professionals from the American Heart Association. Circulation 2017, 135, e927–e999, doi:10.1161/CIR.0000000000000484.

22. Herberg, J.A., Kaforou, M., Gormley, S., Sumner, E.R., Patel, S., Jones, K.D.J., Paulus, S., Fink, C., Martinon-Torres, F., Montana, G., et al. Transcriptomic profiling in childhood H1N1/09 influenza reveals reduced expression of protein synthesis genes. J. Infect. Dis. 2013, 208, 1664–1668, doi:10.1093/infdis/jit348.

23. Ritchie, M.E., Phipson, B., Wu, D., Hu, Y., Law, C.W., Shi, W., Smyth, G.K. limma powers differential expression analyses for RNA-sequencing and microarray studies. Nucleic Acids Res. 2015, 43, e47–e47, doi:10.1093/nar/gkv007.

24. Raudvere, U., Kolberg, L., Kuzmin, I., Arak, T., Adler, P., Peterson, H., Vilo, J. g:Profiler: a web server for functional enrichment analysis and conversions of gene lists (2019 update). Nucleic Acids Res. 2019, 47, W191–W198, doi:10.1093/nar/gkz369.

25. Newman, A.M., Steen, C.B., Liu, C.L., Gentles, A.J., Chaudhuri, A.A., Scherer, F., Khodadoust, M.S., Esfahani, M.S., Luca, B.A., Steiner, D., et al. Determining cell type abundance and expression from bulk tissues with digital cytometry. Nat. Biotechnol. 2019, 37, 773–782, doi:10.1038/s41587-019-0114-2.

26. Charrad, M., Ghazzali, N., Boiteau, V., Niknafs, A. Nbclust: An R package for determining the relevant number of clusters in a data set. J. Stat. Softw. 2014, doi:10.18637/jss.v061.i06.

27. Kimura, Y., Yanagimachi, M., Ino, Y., Aketagawa, M., Matsuo, M., Okayama, A., Shimizu, H., Oba, K., Morioka, I., Imagawa, T., et al. Identification of candidate diagnostic serum biomarkers for Kawasaki disease using proteomic analysis. Sci. Rep. 2017, doi:10.1038/srep43732.

28. Whitin, J.C., Yu, T.T.S., Ling, X.B., Kanegaye, J.T., Burns, J.C., Cohen, H.J. A novel truncated form of serum amyloid a in kawasaki disease. PLoS One 2016, 11, doi:10.1371/journal.pone.0157024.

29. Tibshirani, R. Regression Shrinkage and Selection via the Lasso. J. R. Stat. Soc. Ser. B 1996, 58, 267–288.

30. Danese, S. Nonimmune cells in inflammatory bowel disease: from victim to villain. Trends Immunol. 2008, 29, 555–564.

31. Tanaka, K., Yoshioka, T., Bieberich, C., Jay, G. Role of the Major Histocompatibility Complex Class I Antigens in Tumor Growth and Metastasis. Annu. Rev. Immunol. 1988, 6, 359–380, doi:10.1146/annurev.iy.06.040188.002043.

32. Tremoulet, A.H., Jain, S., Chandrasekar, D., Sun, X., Sato, Y., Burns, J.C. Evolution of laboratory values in patients with Kawasaki disease. Pediatr. Infect. Dis. J. 2011, 30, 1022–1026, doi:10.1097/INF.0b013e31822d4f56.

33. Biezeveld, M.H., van Mierlo, G., Lutter, R., Kuipers, I.M., Dekker, T., Hack, C.E., Newburger, J.W., Kuijpers, T.W. Sustained activation of neutrophils in the course of Kawasaki disease: an association with matrix metalloproteinases. Clin. Exp. Immunol. 2005, 141, 183–188, doi:10.1111/j.1365-2249.2005.02829.x.

34. Asano, T., Ogawa, S. Expression of IL-8 in Kawasaki disease. Clin. Exp. Immunol. 2000, 122, 514–519, doi:10.1046/j.1365-2249.2000.01395.x.

35. Zandstra, J., van de Geer, A., Tanck, M.W.T., van Stijn-Bringas Dimitriades, D., Aarts, C.E.M., Dietz, S.M., van Bruggen, R., Schweintzger, N.A., Zenz, W., Emonts, M., et al. Biomarkers for the Discrimination of Acute Kawasaki Disease From Infections in Childhood. Front. Pediatr. 2020, 8, 355, doi:10.3389/fped.2020.00355.

36. Manlhiot, C., Mueller, B., O’Shea, S., Majeed, H., Bernknopf, B., Labelle, M., Westcott, K. V., Bai, H., Chahal, N., Birken, C.S., et al. Environmental epidemiology of Kawasaki disease: Linking disease etiology, pathogenesis and global distribution. PLoS One 2018, 13, pdoi:10.1371/journal.pone.0191087.

37. Rodó, X., Curcoll, R., Robinson, M., Ballester, J., Burns, J.C., Cayan, D.R., Lipkin, W.I., Williams, B.L., Couto-Rodriguez, M., Nakamura, Y., et al. Tropospheric winds from northeastern China carry the etiologic agent of Kawasaki disease from its source to Japan. Proc. Natl. Acad. Sci. U. S. A. 2014, 111, 7952–7957, doi:10.1073/pnas.1400380111.

38. Lu, X., Zhang, L., Du, H., Zhang, J., Li, Y.Y., Qu, J., Zhang, W., Wang, Y., Bao, S., Li, Y., et al. SARS-CoV-2 Infection in Children. N. Engl. J. Med. 2020, 382, 1663–1665, doi:10.1056/NEJMc2005073.

39. Götzinger, F., Santiago-García, B., Noguera-Julián, A., Lanaspa, M., Lancella, L., Calò Carducci, F.I., Gabrovska, N., Velizarova, S., Prunk, P., Osterman, V., et al. COVID-19 in children and adolescents in Europe: a multinational, multicentre cohort study. Lancet Child Adolesc. Heal. 2020, 0, doi:10.1016/S2352-4642(20)30177-2.

40. Davies, P., Evans, C., Kanthimathinathan, H.K., Lillie, J., Brierley, J., Waters, G., Johnson, M., Griffiths, B., du Pré, P., Mohammad, Z., et al. Intensive care admissions of children with paediatric inflammatory multisystem syndrome temporally associated with SARS-CoV-2 (PIMS-TS) in the UK: a multicentre observational study. Lancet Child Adolesc. Heal. 2020, 0, doi:10.1016/S2352-4642(20)30215-7.

41. Burgner, D., Davila, S., Breunis, W.B., Ng, S.B., Li, Y., Bonnard, C., Ling, L., Wright, V.J., Thalamuthu, A., Odam, M., et al. A genome-wide association study identifies novel and functionally related susceptibility Loci for Kawasaki disease. PLoS Genet 2009, doi:10.1371/journal.pgen.1000319.

42. Curtis, N., Zheng, R., Lamb, J.R., Levin, M. Evidence for a superantigen mediated process in Kawasaki disease. Arch. Dis. Child. 1995, 72, 308–311, doi:10.1136/adc.72.4.308.

43. Han, S.B., Lee, S.Y. Antibiotic use in children with Kawasaki disease. World J. Pediatr. 2018, 14, 621–622.

44. Diz, A.P., Martínez-Fernández, M., Rolán-Alvarez, E. Proteomics in evolutionary ecology: Linking the genotype with the phenotype. Mol. Ecol. 2012, 21, 1060–1080.

45. Benseler, S.M., McCrindle, B.W., Silverman, E.D., Tyrrell, P.N., Wong, J., Yeung, R.S.M. Infections and Kawasaki disease: Implications for coronary artery outcome. Pediatrics 2005, 116, e760–e766, doi:10.1542/peds.2005-0559.

46. Martinón-Torres, F., Salas, A., Rivero-Calle, I., Cebey-López, M., Pardo-Seco, J., Herberg, J.A., Boeddha, N.P., Klobassa, D.S., Secka, F., Paulus, S., et al. Life-threatening infections in children in Europe (the EUCLIDS Project): a prospective cohort study. Lancet Child Adolesc. Heal. 2018, 2, 404–414, doi:10.1016/S2352-4642(18)30113-5.

47. Du, P., Kibbe, W.A., Lin, S.M. lumi: A pipeline for processing Illumina microarray. Bioinformatics 2008, doi:10.1093/bioinformatics/btn224.

48. Leek, J.T., Johnson, W.E., Parker, H.S., Jaffe, A.E., Storey, J.D. The SVA package for removing batch effects and other unwanted variation in high-throughput experiments. Bioinformatics 2012, 28, 882–883, doi:10.1093/bioinformatics/bts034.

49. R Foundation for Statistical Computing R: A language and environment for statistical computing. R A Lang. Environ. Stat. Comput. 3.3.1 2016.

50. Cox, J., Mann, M. MaxQuant enables high peptide identification rates, individualized p.p.b.-range mass accuracies and proteome-wide protein quantification. Nat. Biotechnol. 2008, 26, 1367–1372, doi:10.1038/nbt.1511.

51. Cox, J., Hein, M.Y., Luber, C.A., Paron, I., Nagaraj, N., Mann, M. Accurate proteome-wide label-free quantification by delayed normalization and maximal peptide ratio extraction, termed MaxLFQ. Mol. Cell. Proteomics 2014, 13, 2513–2526, doi:10.1074/mcp.M113.031591.

52. Sweeney, T.E. COCONUT: COmbat CO-Normalization Using conTrols (COCONUT). R package version 1.0.2. 2017.

53. Benjamini, Y., Hochberg, Y. Controlling the False Discovery Rate: A Practical and Powerful Approach to Multiple Testing. J. R. Stat. Soc. Ser. B 1995, 57, 289–300.

54. Supek, F., Bošnjak, M., Škunca, N., Šmuc, T. REVIGO Summarizes and Visualizes Long Lists of Gene Ontology Terms. PLoS One 2011, 6, e21800, doi:10.1371/journal.pone.0021800.

55. Hartigan, J.A., Wong, M.A. Algorithm AS 136: A K-Means Clustering Algorithm. Appl. Stat. 1979, 28, 100, doi:10.2307/2346830.

56. Krzanowski, W.J., Lai, Y.T. A Criterion for Determining the Number of Groups in a Data Set Using Sum-of-Squares Clustering. Biometrics 1988, 44, 23, doi:10.2307/2531893.

57. Caliñski, T., Harabasz, J. A Dendrite Method For Cluster Analysis. Commun. Stat. 1974, 3, 1–27, doi:10.1080/03610927408827101.

58. Gordon, A.D., Hartigan, J.A. Clustering Algorithms. J. Am. Stat. Assoc. 1976, doi:10.2307/2286880.

59. McClain, J.O., Rao, V.R. CLUSTISZ: A Program to Test for the Quality of Clustering of a Set of Objects. J. Mark. Res. 12, 456–460.

60. Dunn, J.C. Well-separated clusters and optimal fuzzy partitions. J. Cybern. 1974, 4, 95–104, doi:10.1080/01969727408546059.

61. Halkidi, M., Vazirgiannis, M., Balislakis, V. Quality scheme assessment in the clustering process. In Proceedings of the Lecture Notes in Computer Science (including subseries Lecture Notes in Artificial Intelligence and Lecture Notes in Bioinformatics); Springer Verlag, 2000; Vol. 1910, pp. 265–276.

62. Halkidi, M., Vazirgiannis, M. Clustering validity assessment: Finding the optimal partitioning of a data set. In Proceedings of the Proceedings - IEEE International Conference on Data Mining, ICDM; 2001; pp. 187–194.

63. Hubert, L.J., Levin, J.R. A general statistical framework for assessing categorical clustering in free recall. Psychol. Bull. 1976, 83, 1072–1080, doi:10.1037/0033-2909.83.6.1072.

64. Rousseeuw, P.J. Silhouettes: A graphical aid to the interpretation and validation of cluster analysis. J. Comput. Appl. Math. 1987, 20, 53–65, doi:10.1016/0377-0427(87)90125-7.

65. Ball, G.H., Hall, D.J. ISODATA, A NOVEL METHOD OF DATA ANALYSIS AND PATTERN CLASSIFICATION. undefined 1965.

66. Milligan, G.W. An examination of the effect of six types of error perturbation on fifteen clustering algorithms. Psychometrika 1980, 45, 325–342, doi:10.1007/BF02293907.

67. Milligan, G.W. A monte carlo study of thirty internal criterion measures for cluster analysis. Psychometrika 1981, 46, 187–199, doi:10.1007/BF02293899.

68. Ratkowsky, D., Lance, G. A Criterion for Determining the Number of Groups in a Classification. Aust. Comput. J. 1978, 10, 115–117.

69. Robin, X., Turck, N., Hainard, A., Tiberti, N., Lisacek, F., Sanchez, J.C., Müller, M. pROC: An open-source package for R and S+ to analyze and compare ROC curves. BMC Bioinformatics 2011, 12, 77, doi:10.1186/1471-2105-12-77.

